# Basal UPR activity in *Aspergillus fumigatus* regulates adaptation to nutrient stress and is critical for the establishment of corneal infection

**DOI:** 10.1101/2023.05.22.541860

**Authors:** Manali M. Kamath, Jorge D. Lightfoot, Emily M. Adams, Becca L. Wells, Kevin K. Fuller

**Author notes:** Correspondence: Kevin K. Fuller, PhD., Assistant Professor, 608 Stanton L. Young Dr., Oklahoma City, OK, 73104.

## Abstract

The *Aspergillus fumigatus* unfolded protein response (UPR) is a two-component relay consisting of the ER-bound IreA protein, which splices and activates the mRNA of the transcription factor HacA. Spliced *hacA* accumulates under conditions of acute ER stress *in vitro*, and UPR null mutants are hypovirulent in a murine model of invasive pulmonary infection. In this report, we demonstrate that a *hacA* deletion mutant is completely unable to establish infection in a model of fungal keratitis, a corneal infection and an important cause of ocular morbidity and unilateral blindness worldwide. Contrary to our initial prediction, however, we demonstrate that *hacA* splicing is not increased above baseline conditions in the cornea, nor is the expression of genes classically associated with UPR activation, such as protein chaperones. We employed transcriptomics on wild-type and *ΔhacA* strains in gelatin media, as a proxy for the corneal environment, and found that *hacA* supports the expression of numerous primary and secondary metabolic processes that likely promote adaptation to nutrient limitation. Taken together, our results support a model in which the cornea, similar to growth on protein *in vitro*, is a source of sub-acute ER stress for *A. fumigatus*, but one nevertheless that requires the UPR pathway for proper adaptation. The data also suggest that this pathway could be a target for novel antifungals that improve visual outcomes for fungal keratitis patients.

**AUTHOR SUMMARY:** Fungal keratitis has emerged as a leading cause of ocular morbidity and unilateral blindness worldwide. Relative to other infectious contexts, however, little is known about the fungal genes or pathways that regulate invasive growth and virulence in the corneal environment. In this report, we demonstrate that genetic disruption of the *Aspergillus fumigatus* unfolded protein response (UPR) abolishes the ability of the mold to establish infection in a mouse model of FK. Despite this critical role for virulence, however, we did not detect a concerted activation of the pathway beyond levels observed on standard medium, suggesting that the host environment is not an acute source of endoplasmic reticulum stress. Transcriptomic profiling of the wild-type and UPR-deficient strains under host-relevant nutrient conditions revealed a critical role for the pathway in regulating primary and secondary metabolism, cell wall biology, and mitochondrial function, all of which likely modulate fungal growth within and interactions with the host. These results expand our understanding of UPR regulation and function in this important mold pathogen and suggest the pathway could serve as a target for novel antifungals that improve visual outcomes in the setting of fungal keratitis.

## INTRODUCTION

The pathogenic potential of *Aspergillus fumigatus* and other fungi is tied to the functional integrity of the endoplasmic reticulum (ER). First, the organelle serves as the synthetic hub for a myriad of cell surface or secreted virulence proteins, including 1) cell wall remodeling enzymes that facilitate apical extension and stress resistance, 2) hydrolases that promote nutrient acquisition from, and penetration through host tissue, and 3) transporters involved in macro/micronutrient uptake or efflux (1–5). Second, numerous stresses encountered within the host environment e.g., hypoxia, oxidative stress, or nutrient limitation that places a demand on hydrolase secretion may cumulatively lead to a cytotoxic aggregation of unfolded peptides in the ER that will induce cell death if left unchecked (2,6,7). The unfolded protein response (UPR) is an evolutionarily conserved signaling pathway that senses ER stress and drives a transcriptional response that restores protein folding and secretory homeostasis (2,8,9). Thus, by virtue of its role in ER function, the UPR is a de facto regulator of virulence and a putative target for novel antifungal therapy.

The UPR of *A. fumigatus* is a two-component relay that generally follows the canonical pathway described in *Saccharomyces cerevisiae* (10–12). The sensing module, IreA, is an ER transmembrane protein whose luminal domain interacts with the protein chaperone BipA/Kar2 under homeostatic conditions. BipA dissociates upon the accumulation of unfolded proteins in the lumen, leading to an oligomerization of IreA molecules that promotes the trans-autophosphorylation of its cytosolic kinase domain and activation of its endoribonuclease domain. The latter splices a 20 bp unconventional intron from the cytosolic *hacA* transcript (*hacA*^u^, <u>u</u>ninduced), leading to a frame-shifted isoform (*hacA*^i^, induced) that encodes a bZIP transcription factor that provides output for the UPR (13,14). Work from Askew and colleagues demonstrated that 1) *hacA^i^* is the most abundant splice form in *A. fumigatus* following treatment with the thiol-reducing agent dithiothreitol (DTT) and, 2) treatment with DTT or tunicamycin, which inhibits protein glycosylation and normal peptide folding, leads to the induction of ∼60 genes in an IreA and HacA dependent manner (15). This “inducible UPR” (iUPR) consists primarily of genes that promote protein folding (e.g. chaperones, foldases, protein glycosylases) and vesicle trafficking, which is consistent with known UPR-regulated genes in other fungi (15). The *A. fumigatus* UPR also regulates the expression of calcium ATPases that facilitate the influx of Ca2+ ions needed for chaperone function (16). It follows, therefore, that both *ireA* or *hacA* deletion mutants are hypersensitive to stressors that acutely unfold proteins (high temperature, DTT, TM) or block the egress of proteins from the ER (brefeldin A) (16–18). The *ΔhacA* or *ΔireA* strains are furthermore growth defective on protein-rich media, such as skim milk agar, which correlates with a lack of detectable collagenase activity from culture supernatants (14). This suggests that the enhanced secretory burden experienced on polymeric substrates is another source of ER stress that promotes increased *hacA* splicing and UPR induction.

Although *hacA^u^* is the dominant splice form under homeostatic conditions (e.g. rich media), there is nevertheless detectable *hacA^i^* product at all times. This tracks with a slight growth and conidiation defect of the *ΔhacA* or *ΔireA* strains on standard lab media and suggests that HacA buffers low-level ER stress associated with normal growth and developmental processes. Interestingly, the microarray study by Feng et al. identified ∼250 genes that are regulated by IreA and HacA on a rich medium, only 9 of which overlapped with the iUPR defined after DTT and TM treatment (15). This so-called “basal UPR” (bUPR) includes genes enriched in oxidative metabolism and mitochondrial function, which is distinct from the enriched gene categories associated with the iUPR. One possible explanation is that the bUPR and iUPR are two ends of an activity spectrum that is fine-tuned at the level of *hacA* splicing. In this scenario, UPR gene expression is solely dependent on the canonical HacA protein encoded by the *hacA^i^* mRNA. Another possibility is that the bUPR is qualitatively distinct and driven by a protein product encoded by the *hacA^u^* mRNA (19). To date, the UPR-dependent transcriptome of *A. fumigatus* has only been evaluated under conditions of low (rich medium or acute (e.g. DTT)) ER stress, and so a more intermediate level UPR activation and output, if it exists, has yet to be interrogated.

The importance of the UPR in *A. fumigatus* pathogenesis has been evaluated in a murine model of invasive pulmonary aspergillosis (IPA), where *ΔhacA* displays an approximately 50% reduction in virulence based on cumulative mortality (14). The role of HacA in this regard, as eluded to above, is likely multifactorial. Individual phenotypes associated with *ΔhacA* including reduced hydrolase secretion, a hypersensitivity to iron limitation and cell wall stress each have a presumed or demonstrated role in fungal adaptation to the lung environment and probably contribute additively to the virulence phenotype. Interestingly, the *ΔireA* mutant was completely avirulent in the same infection model. This is consistent with the fact that *ΔireA* displays a greater growth defect at 37°C and an increased sensitivity to iron and nutrient limitation (relative to *ΔhacA*). While this supports a model in which IreA has yet-to-be identified functions beyond *hacA* splicing, the virulence of *ΔireA* could be restored to wild-type levels by introducing a constitutively spliced *hacA* (*hacAi*) allele (15). Nevertheless, the relationship between the iUPR and bUPR in virulence remains incompletely understood. While it may be assumed that a myriad of host-derived stresses leads to acute ER stress and activation of iUPR, the demonstration of *hacA* splicing *in vivo* has not been formally examined.

Beyond the lung, *A. fumigatus* is also an important causative agent of fungal keratitis (FK), a sight-threatening infection of the cornea that affects 1-2 million individuals globally (20). The disease occurs when fungal spores or hyphal fragments gain entry to the corneal tissue following damage to the protective epithelium, predominantly as a result of agriculture-related trauma or contact lens wear (21,22). In all contexts, however, the combination of invasive hyphal growth and leukocytic infiltration causes acute pain, photophobia, and a disruption to the optical properties of the cornea that result in vision loss (23–25). FK results in the need for corneal transplantation in ∼40% of cases and a wholesale removal of the eye in about 10% (20). As with IPA, the poor outcomes of FK is due, in part, to the inadequacy of current antifungals, for which only the polyene natamycin has received FDA-approved for corneal use (26,27). Thus, it is a major goal of our group to identify *A. fumigatus* pathways that support virulence in both the lung and corneal environments, allowing for the development of novel antifungals with dual use. The primary site of fungal growth during FK is the corneal stroma, which is avascular and composed primarily of a dense collagen matrix (28). We reasoned that growth and penetration in this environment would require upregulation of secreted collagenases that would induce ER stress, lead to increased *hacA* splicing and ultimately render the UPR critical for virulence. In this report, we do indeed demonstrate that a *ΔhacA* mutant is unable to establish infection in a murine model of FK. Interestingly, however, analysis of fungal RNA from infected corneas did not reveal increased *hacA* splicing beyond that observed under baseline *in vitro* culture. To gain a deeper insight into how the UPR might regulate adaptation and growth within an environment that resembles the host, we performed RNA-sequencing in wild-type versus *ΔhacA* strains following culture in media containing glucose or gelatin as the sole carbon source. In the wild-type, gelatin media did not promote the upregulation of genes typically associated with the iUPR, suggesting nutrient limitation alone is not sufficient to acutely activate the canonical IreA-HacA pathway. Nevertheless, the pathway does regulate key metabolic pathways involved in the growth on gelatin, including those involved in alternative carbon utilization, cell wall homeostasis, redox balance, and secondary metabolism. These data suggest that the cornea, and perhaps other host environments, fail to drive an iUPR response and instead lead to an intermediate level of UPR activation that has a unique transcriptional signature. Moreover, the data suggest that the pathway could serve as a target for novel drugs that improve patient outcomes in the setting of FK.

## RESULTS

### *A. fumigatus* protease-encoding genes are upregulated in a murine model of FK

We first reasoned that *A. fumigatus* upregulates the expression of various secreted proteases during corneal infection, either to mediate penetration through the dense collagen matrix or for nutritional support within an environment that is ostensibly poor in preferred carbon or nitrogen substrates. To test this, we employed an epithelial abrasion model of FK developed and previously reported by our group (20,29). Briefly, C57BL/6J mice were immunosuppressed with a single treatment of methylprednisolone on the day preceding inoculation (**Fig S1**). Although immunosuppression is not a prerequisite for the establishment of FK in patients or our model, we have found that a single dose of steroids facilitates more uniform disease progression across animals. On the day of inoculation, the epithelium on the cornea was ulcerated and overlayed with *A. fumigatus* (Af293) conidia that were pre-germinated (swollen, but not polarized) in rich media. In contrast to the sham-inoculated controls, *A. fumigatus* corneas showed obvious signs of infection by 48 h post-inoculation (p.i.), including diffuse corneal opacification and surface irregularity (**Fig 1A**). Total RNA was isolated from infected corneas at this time point as well as from *A. fumigatus* cultured in glucose minimal medium (GMM), containing a preferred source of carbon (glucose) and nitrogen (NH_4_), for 48h as the baseline condition. Consistent with our hypothesis, the corneal-derived samples displayed higher levels (50-1000 fold) of steady-state mRNA for each of *A. fumigatus* collagenase genes tested, including the metalloprotease (*mep1* – AFUA_1G07730), a serine alkaline protease (*alp1* – AFUA_4G11800), and two dipeptidyl-peptidases (*dppIV* – AFUA_4G09320 and *dppV –* AFUA_2G09030) (**Fig 1B**). These results support the interpretation that *A. fumigatus* upregulates its secreted hydrolytic activity to support invasive growth in the corneal environment.

**Fig 1.**
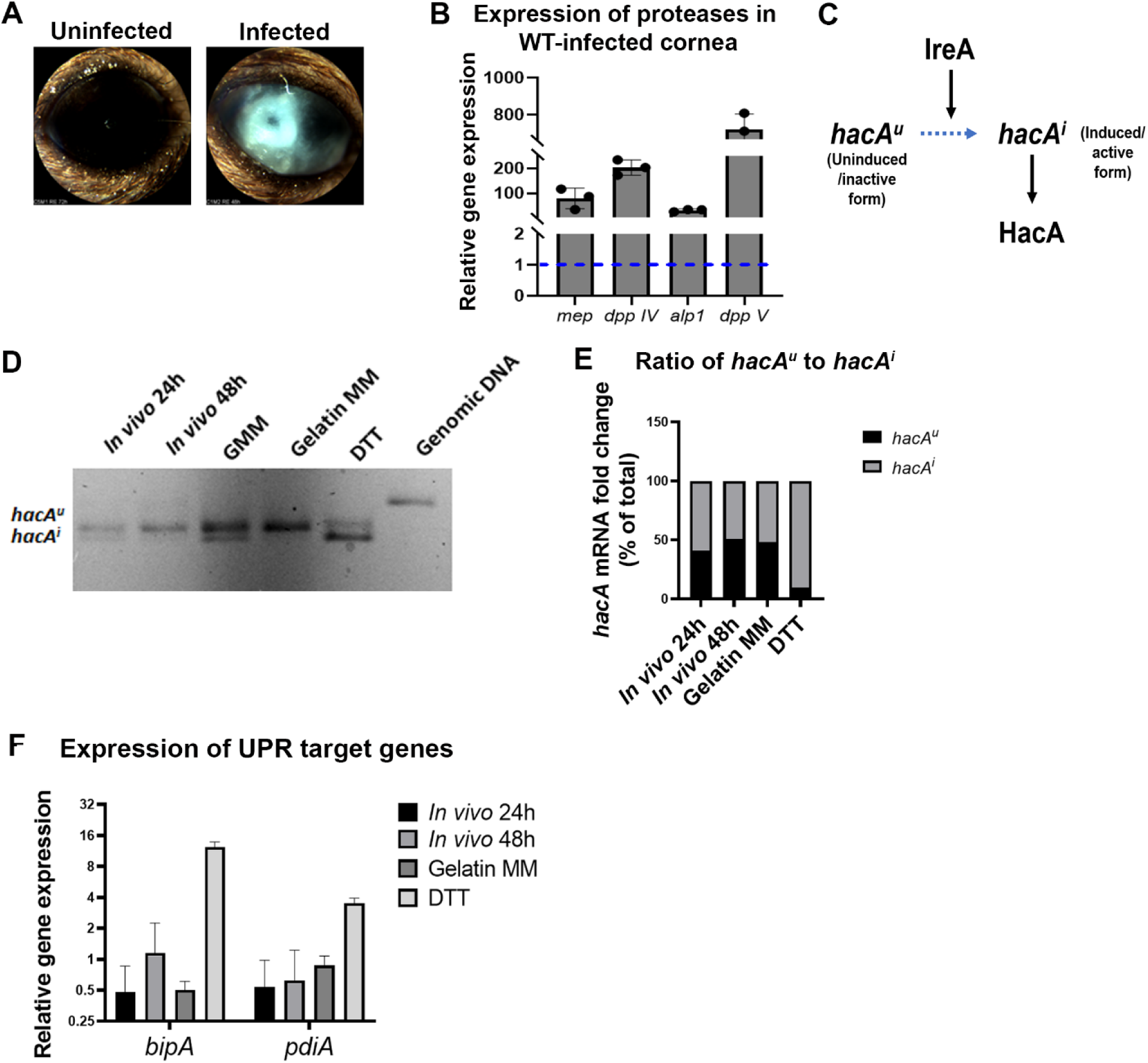
UPR activation occurs at a basal level in *A. fumigatus*. (A) Typical corneal images 48 h post-infection of either sham (PBS) or WT *A. fumigatus*-infected murine eyes using the epithelial abrasion model of keratitis, (B) qRT-PCR revealed an up-regulation of protease genes with collagenase activity from WT *A. fumigatus*-infected cornea normalized to β-tubulin, data represent the mean of triplicate samples, (C) Schematic diagram illustrating the splicing of the uninduced form of *hacA* (*hacA^u^*) to form the induced form (*hacA^i^*) which then forms the HacA functional protein, (D) The *hacA* mRNA was amplified by RT-PCR and the PCR products were separated on a 3% agarose gel to detect both the *hacA^u^* (upper band) and *hacA^i^* (lower band) forms in the *in vivo* (24 and 48h p.i.) and the *in vitro* (GMM, gelatin MM, and DTT-treated) samples. qRT-PCR depicting (E) the ratio of *hacA^i^* to *hacA^u^* mRNA and (F) expression of UPR target genes – *bipA* and *pdiA* in the WT *A. fumigatus-*infected cornea at 24 h & 48 h p.i., gelatin MM and GMM treated with DTT relative to the control (GMM).

### The Af293 UPR is not induced beyond basal levels during nutrient limitation or corneal infection

It is known that treatment of *A. fumigatus* with dithiothreitol (DTT) leads to a rapid accumulation of misfolded proteins in the ER and the IreA-mediated splicing of the *hacA* transcript from the uninduced (*hacA*^u^) to the induced/active (*hacA*^i^) form (**Fig 1C**). We predicted that the increased expression of secreted proteases that occurs in the corneal environment would similarly promote ER stress and a detectable accumulation of *hacA*^i^ beyond baseline conditions. To test this, we analyzed the relative abundance of *hacA* splice forms in *A. fumigatus* Af293 grown in the following conditions: (1) GMM, as the baseline (non-inducing) control; (2) GMM with 1 h DTT treatment as a positive control; (3) gelatin minimal medium (Gel MM), in which protease gene expression is induced similar to the cornea and; (4) 48 h murine corneas as described above. We performed RT-PCR using *hacA* primers that span the previously defined 20 bp non-conventional splice region, thus allowing for the detection of both the *hacA^u^* and *hacA^i^* forms following the resolution of the amplicons by gel electrophoresis. The samples were also analyzed by qRT-PCR using two primer sets that distinctly amplify the *hacA^u^* and *hacA^i^* cDNAs. Both forms of the *hacA* band were detected in the baseline sample, although the un-induced (higher molecular weight) band was more abundant. We interpret this steady state *hacA* splicing to correspond with the ‘basal UPR’ (bUPR) previously described for *A. fumigatus* strain AfS28 (15). The 1h DTT treatment effectively reversed the relative intensity of the two bands, which is again consistent with previous reports and represents the presence of the ‘induced UPR’ (iUPR) (**Fig 1D and 1E**). In contrast to our prediction, however, neither growth in gelatin nor the cornea shifted the splice form ratio beyond what was observed in GMM. The *hacA* splicing data corresponded with the expression analysis of two known genes under the control of the HacA transcription factor, the protein chaperones *bipA* and *pdiA*, which were both induced in DTT but not in gelatin or the corneal samples (**Fig 1F**).

### Transcriptomics supports a distinct role for the UPR under acute ER and nutritional stresses

To further interrogate whether ER/UPR stress is induced in proteinaceous environments, we analyzed the above-described gelatin and DTT samples by RNA sequencing. Differentially expressed genes (DEGs) were defined as those with a 4-fold change relative to the GMM condition. DTT treatment led to the induction of 345 genes, including enrichment of those within the ‘protein folding and stabilization’ Functional (FunCat) categories. Such genes including protein chaperones (*bipA*, *hsp70*), calnexin, protein isomerases, and ER resident oxidoreductases and ATPases were similarly found to be induced in the *A. fumigatus* AfS28 background by microarray, and therefore appear to constitute a conserved iUPR (14). Other enriched FunCat terms in our DTT-induced gene set included ‘detoxification by modification’, ‘secondary metabolism’, ‘oxidative stress’, and ‘extracellular polysaccharide degradation’, suggesting a broad stress response that may or may not be attributed to UPR signaling (**Table 1**).

**Table 1:** Term enrichment analysis of Af293 DEGs under ER stress conditions.

|  | FunCat ID | FunCat description | Exact p-value | # genes/input |
| --- | --- | --- | --- | --- |
| Gelatin<br>MM | 1.2 | Secondary metabolism | 8.46436e-20 | 122 / 672 |
|  | 20.09.18.07 | Non-vesicular cellular import | 6.706738e-8 | 33 / 672 |
|  | 1.05 | C-compound and carbohydrate metabolism | 2.559836e-7 | 90 / 672 |
|  | 2.25 | Oxidation of fatty acids | 3.239912e-7 | 16 / 672 |
|  | 16.21.01 | Heme binding | 1.39322E-06 | 16 / 672 |
|  | 01.06.05 | Fatty acid metabolism | 2.16031E-06 | 25 / 672 |
|  | 01.01.11.04.02 | Degradation of leucine | 2.1768E-06 | 9 / 672 |
|  | 20.01.03 | C-compound and carbohydrate transport | 2.92668E-06 | 34 / 672 |
|  | 01.01.09.04.02 | Degradation of phenylalanine | 5.82589E-06 | 11 / 672 |
|  | 20.03 | Transport facilities | 7.16462E-06 | 52 / 672 |
|  | 20.09.18 | Cellular import | 7.99918E-06 | 36 / 672 |
|  | 16.21.15 | Biotin binding | 1.03532E-05 | 6 / 672 |
|  | 1.06 | Lipid, fatty acid and isoprenoid metabolism | 2.68213E-05 | 61 / 672 |
|  | 20.01.23 | Allantoin and allantoate transport | 5.68402E-05 | 12 / 672 |
|  | 01.01.09.05.02 | Degradation of tyrosine | 6.59254E-05 | 7 / 672 |
|  | 01.05.03 | Polysaccharide metabolism | 0.000150225 | 30 / 672 |
|  | 2.09 | Anaplerotic reactions | 0.000250594 | 3 / 672 |
|  | 20.01.15 | Electron transport | 0.000549018 | 34 / 672 |
|  | 32.07.01 | Detoxification involving cytochrome P450 | 0.001301791 | 10 / 672 |
|  | 16.21.05 | FAD/FMN binding | 0.001655444 | 16 / 672 |
|  | 32.07.07.01 | Catalase reaction | 0.002276066 | 3 / 672 |
|  | 01.20.33 | Metabolism of secondary products derived from L-tryptophan | 0.002397907 | 5 / 672 |
| DTT | 1.2 | Secondary metabolism | 3.015158e-10 | 50 / 345 |
|  | 32.07.03 | Detoxification by modification | 4.78508E-05 | 7 / 345 |
|  | 32.05.01 | Resistance proteins | 0.000174247 | 9 / 345 |
|  | 32.01.01 | Oxidative stress response | 0.000402305 | 9 / 345 |
|  | 14.01 | Protein folding and stabilization | 0.001053691 | 10 / 345 |
|  | 01.25.01 | Extracellular polysaccharide degradation | 0.001258999 | 5 / 345 |

Growth in gelatin minimal medium led to the upregulation of 672 genes (relative to GMM) with term enrichments that indicated a broad shift towards the metabolism of protein and other polymeric substrates, such as fatty acids and polysaccharides. Despite this apparent dependency on secretory activity, however, there was not a corresponding increase in genes involved in protein folding or ER homeostasis observed in DTT. Indeed, only 110 overlapped between the two and included two enriched categories: ‘secondary metabolism’ and polysaccharide binding’ (**Fig 2**).

**Fig 2.**
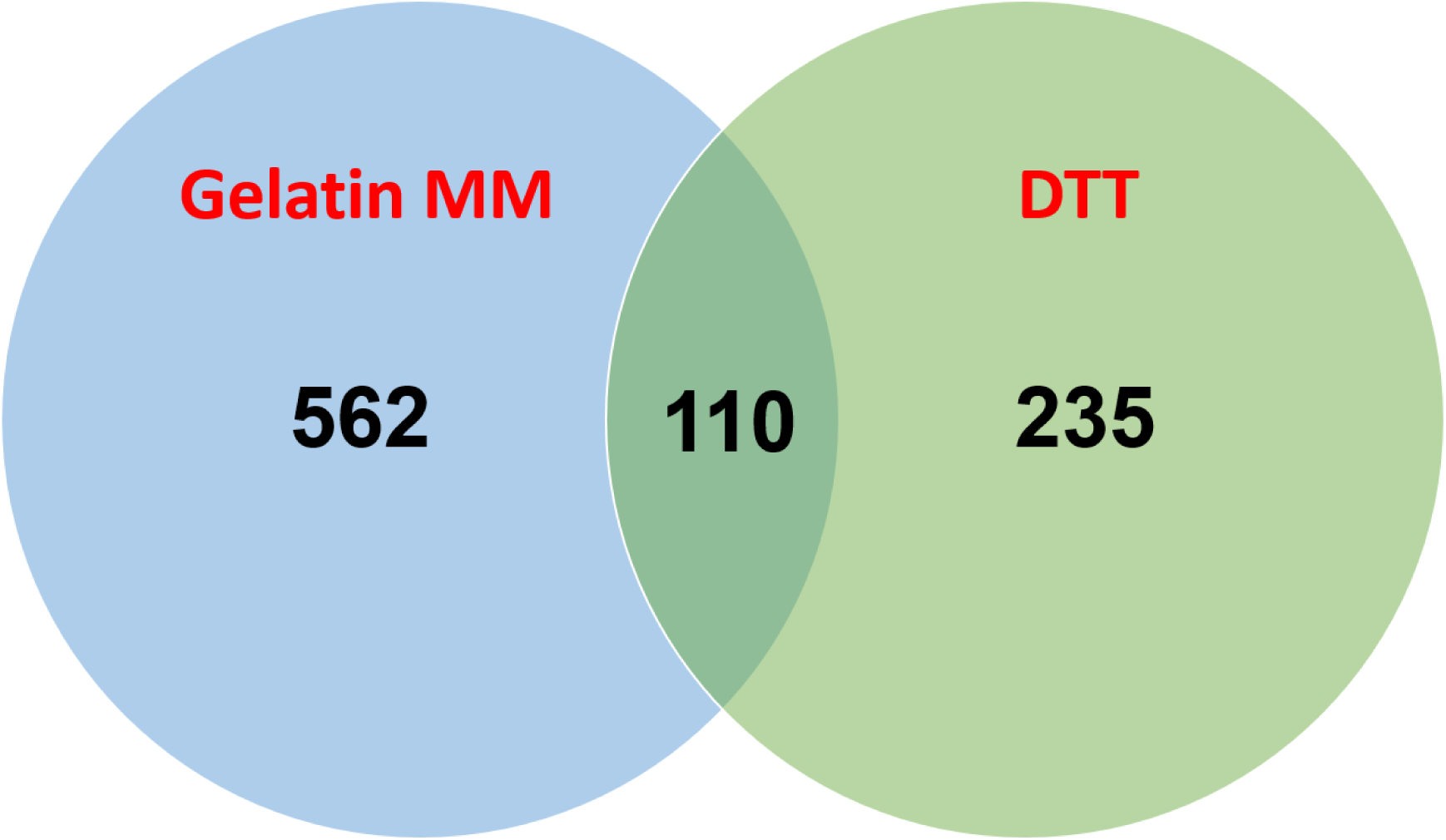
Venn diagram displaying the relationship of DEGs between the two comparisons – Gelatin MM and DTT with a cutoff of fold change > 4.0.

Interestingly, the sensitivity of Af293 to brefeldin A, which induces ER stress by blocking vesicular transport through the Golgi, was increased on gelatin compared to GMM (**Fig S2C**). Taken together with the *hacA* splicing and transcriptomics data, it appears that the fungus is under some degree of stress in protein-rich environments, just not to the threshold required for iUPR activation. While we were unable to perform RNA-seq on our corneal samples due to a lack of fungal RNA yield, the lack of both *hacA* splicing and candidate gene induction (e.g. *bipA*) suggests the iUPR is also not active during corneal infection.

In addition to the upregulation of genes encoding secreted hydrolases, we further observed growth on gelatin led to an upregulation of genes enriched for amino acid transport, gluconeogenesis, oxidative metabolism (electron transport), and oxidation-reduction, all of which seemingly corresponded to a broader metabolic shift to alternative carbon (non-glucose) utilization (**Table 1**). Surprisingly, however, we also observed an upregulation of genes involved in iron/heme binding as well as several secondary metabolic clusters, including gliotoxin. While these genes likely do not promote growth in gelatin MM directly, they may be broadly de-repressed as a part of the adaptive response within nutrient-limiting environments that may be concomitantly stressful in other ways. A full list of DEGs and term enrichment analysis in the DTT and gelatin conditions can be found in supplemental **Tables S1 and S2**.

### HacA is required for the growth of Af293 under acute ER stress and on protein-rich substrates

Richie et al. previously demonstrated that the *A. fumigatus* AfS28 *hacA* deletion mutant is impaired for growth on protein-rich substrates, presumptively because the iUPR is operative in such environments. Given our above results in Af293, which suggest only the bUPR is active in gelatin, we wondered to what degree the UPR would regulate growth in this *A. fumigatus* strain. Accordingly, the Af293 *hacA* coding sequence was replaced with the hygromycin resistance cassette (*hph*) using Cas9-mediated homologous recombination (Al Abdallah 2017; Lightfoot and Fuller 2019). Subsequent generation of the complemented strain (*ΔhacA*::*hacA*) was achieved through the ectopic integration of the WT *hacA* allele (fused with the *bleR* cassette) into the *ΔhacA* mutant (**Fig S2A**). Consistent with the AfS28 Δ*hacA* and orthologous mutants in other fungi, the Af293 *ΔhacA* was hypersensitive to stressors that acutely misfold proteins (high temperature), inhibit the normal folding of nascent polypeptides in the ER (tunicamycin), or block the egress of proteins from the ER (brefeldin A) (**Figs 3A, 3B, S2C**). Taken together, these results demonstrate a conserved essentiality of the UPR under conditions of acute ER stress in which the iUPR is functional. Interestingly, the *ΔhacA* mutant displayed minor but obvious growth and conidiation defects under control/baseline conditions (YPD or GMM at 35^0^C), which were fully restored to WT levels in the complemented strain (**Figs 3A, 3E, S2B**). This supports the interpretation that bUPR buffers the endogenous ER stress associated with growth and developmental processes, even in otherwise favorable conditions.

**Fig 3.**
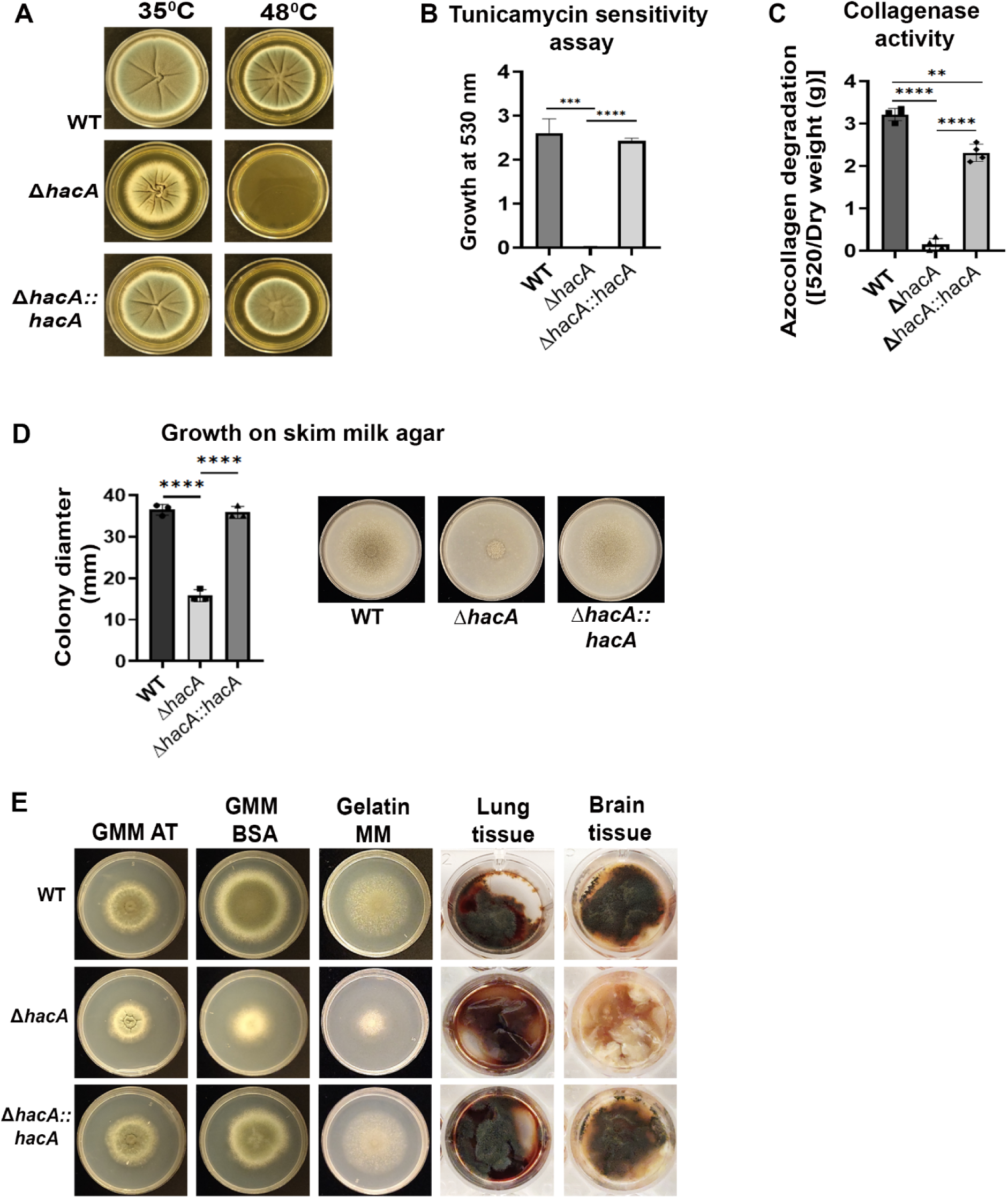
The Δ*hacA* mutant is defective in collagenase secretion and growth on complex substrates. Δ*hacA* mutant is hypersensitive to (A) higher temperature – 48^0^C on YPD agar plates and (B) glycosylation inhibitor – Tunicamycin in liquid GMM-AT; Data represent the mean of triplicate samples analyzed by Brown-Forsythe and Welch’s ANOVA **** <0.0001, ** 0.0022, (C) Culture supernatants were analyzed for azocollagen degradation [absorbance at 520nm normalized to dry weight (g)] after growth in GMM-FBS at 72h; Data represent the mean of four replicate samples analyzed by Brown-Forsythe and Welch’s ANOVA **** <0.0001, ** 0.0022 (D) Representative images of WT, Δ*hacA* and Δ*hacA::hacA* grown on 1% skim milk agar with growth depicted as colony diameter; Data represented as a mean of triplicate samples analyzed by One-way Ordinary ANOVA **** <0.0001. (E) Representative images of growth of the above mentioned strains on media with different protein sources – GMM+BSA, gelatin minimal media as well as explanted lung and brain tissue from mice; Data represented as a mean of triplicate

To determine if the Af293 UPR supports growth in proteinaceous environments, we began by collecting fungal supernatants following growth in minimal media containing albumin as the sole nitrogen source (GMM-BSA). In contrast to the WT and complement-derived samples, *ΔhacA* supernatants displayed undetectable azocollagen degradation, suggesting that the secretion of collagenases was impaired in the mutant (**Fig 3C**). Consistent with this, the Δ*hacA* mutant displayed a partial but significant growth defect on various protein-rich substrates (including GMM-BSA, gelatin minimal medium, and skim milk agar) relative to the WT and *ΔhacA*::*hacA* stains (**Figs 3D, 3E**). The zone of clearance around the *ΔhacA* colony on the skim milk plates indicated residual protease activity that could at least partially account for the growth on these media. More strikingly, *ΔhacA* completely failed to grow on mouse lung and brain tissues, suggesting an increased need for UPR activity on biological substrates (**Fig 3E**). In summary, these data demonstrate that although the UPR pathway is not demonstrably induced in nutrient-limiting environments, it nevertheless plays a role in regulating the secretion of proteases and growth in such environments.

### HacA regulates primary and secondary metabolic shifts associated with growth in gelatin

To gain a more comprehensive insight into how HacA regulates *A. fumigatus* growth in various nutrient environments, we compared the transcriptomes of the Af293 WT and *ΔhacA* strains in GMM and gelatin-MM (**Fig S3A**). In GMM, we found 514 genes were downregulated in *ΔhacA* (relative to WT), including numerous chaperones that fit with the known function of the UPR (e.g. *bipA, hsp78, hsp98, hsp30*) (**Fig 4**). Also downregulated in *ΔhacA* were numerous genes encoding proteins destined for the plasma membrane or secretion, including nutrient and drug transporters, proteases, and polysaccharide binding and degrading enzymes. Beyond the breakdown of polysaccharides as a nutrient source, it is likely many of such genes are involved in the routine maintenance of cell wall (e.g. glucanases, chitinases), and this corresponded to hypersensitivity of *ΔhacA* to cell wall stressors such as Congo red (**Fig S4E**). Thus, during growth in GMM, basal HacA function appears to not only support the expression of genes that regulate ER homeostasis and secretory function but also broadly regulates the expression of genes that traffic through the secretory pathway. The dysregulation of these genes in *ΔhacA* likely accounts for the growth and developmental defects observed on GMM or rich media (**Fig 1**). Interestingly, 328 genes were upregulated in *ΔhacA* relative to the WT, and these were enriched in various secondary metabolic pathways (fumagillin, fumigaclavine C, fumiquinazoline), iron and heme binding, and oxidoreductase activity (**Fig 4B**). Although we predict that the expression of these pathways would not impact growth on GMM per se, their de-repression may be energetically unfavorable and further contribute to the growth defects of *ΔhacA*.

**Fig 4:**
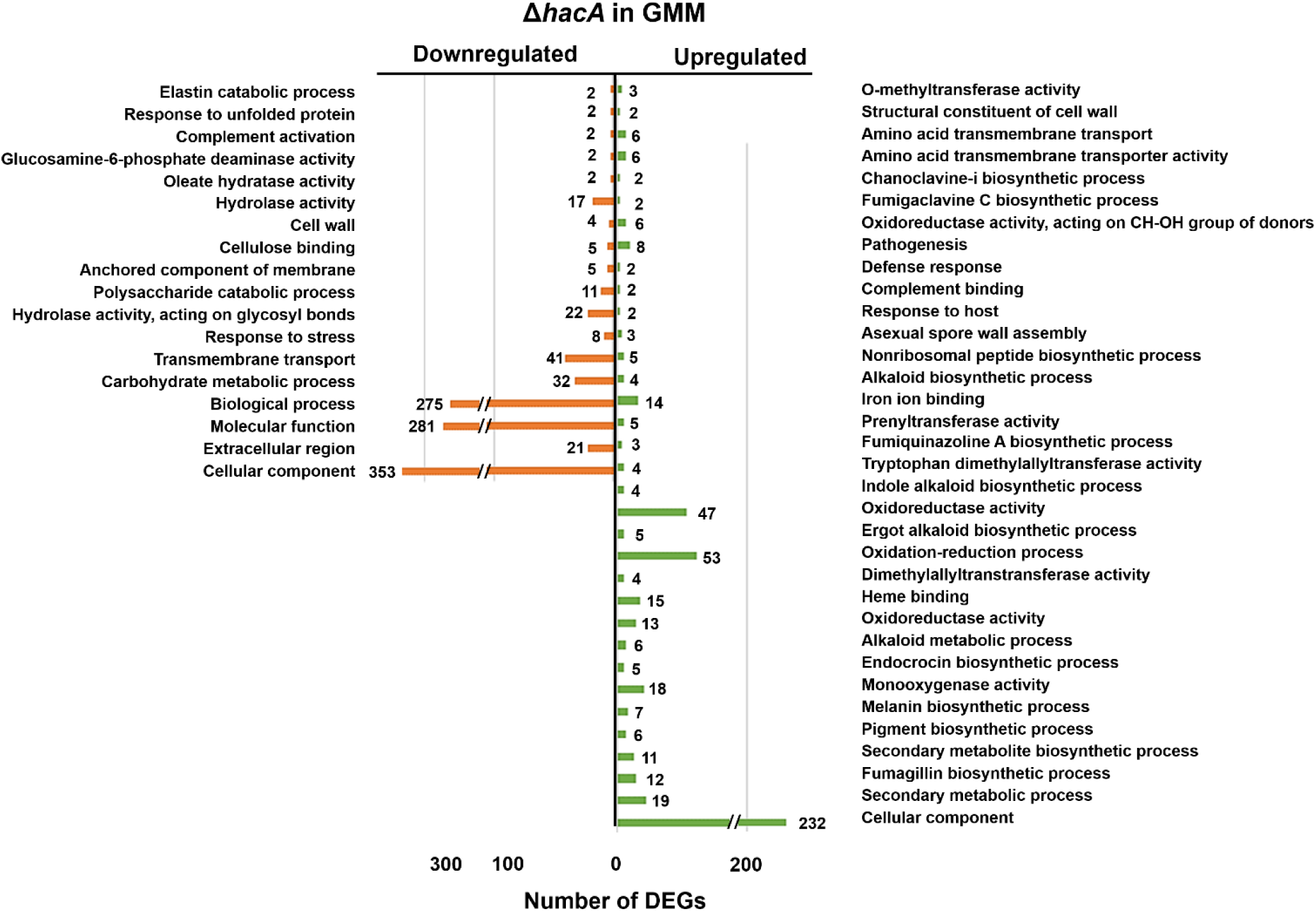
Significantly enriched categories indicating the total number of differentially regulated genes from Gene Ontology (GO) in Δ*hacA* grown in GMM compared to WT are listed with a cut-off of 4-fold change.

In gelatin MM, 311 genes were downregulated in *ΔhacA* relative to WT whereas several of the enriched Gene Ontology (GO) terms for these DEGS were similarly observed among the GMM (e.g., hydrolases, polysaccharide binding/cell wall, hsp/chaperones), the majority were unique to gelatin MM, including oxidation-reduction, iron binding, secondary metabolism, and pigment/melanin biosynthesis categories (**Fig 5A**). Many of these gene categories were upregulated in the WT Af293 in gelatin (relative to WT in GMM), suggesting that HacA plays an important role in regulating the transcriptional adaptation of *A. fumigatus* to the nutrient environment. The dysregulation of several genes from the RNA-seq dataset was validated in a replicated experiment using qRT-PCR (**Fig 5B**). 89 genes were upregulated in *ΔhacA* relative to WT, and these were enriched in just a few FunCat terms, including peptide transport and extracellular polysaccharide degradation. A full list of DEGs and term enrichment analysis between WT and *ΔhacA* in GMM and gelatin can be found in **Table S3 and S4**.

**Fig 5.**
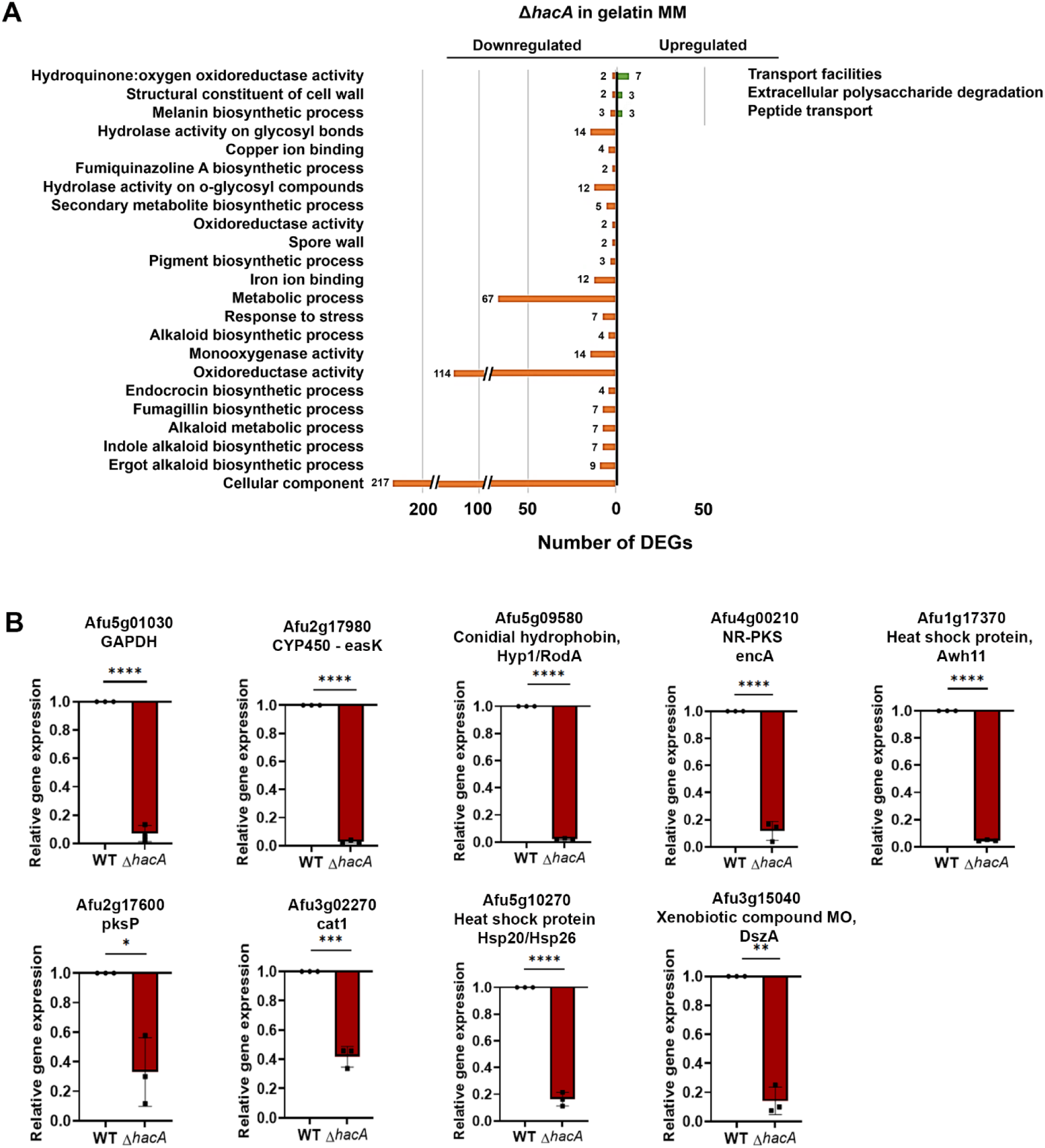
Role of *hacA* under nutritional limitation – gelatin MM. (A) Significantly enriched categories indicating the total number of DEGs under each category from GO (underrepresented) and FunCat (enriched) are listed with a cut-off of 4-fold change. (B) qRT-PCR validation of the RNA-sequencing data using selected genes representing the various categories normalized to actin from WT samples. Data represent a mean of triplicate samples analyzed using the unpaired T-test ** 0.0076, *** 0.0001, **** <0.0001

### UPR is critical in the establishment of the corneal pathogenesis of *A. fumigatus*

Despite the important role HacA plays in secretion and the transcriptional profile in gelatin (collagen) media, the *ΔhacA* mutant nevertheless retains some growth in this environment (**Fig 3**). We, therefore, wondered to what degree, if any, the mutant might be attenuated for growth/virulence in the cornea, in which we predict collagen serves as the major nutritional source. To test this, the Af293 WT, Δ*hacA,* and Δ*hacA::hacA* strains were analyzed in the murine model described above. Remarkably, and in contrast to the progressive disease development that occurred with the WT and complement infected corneas, animals inoculated with Δ*hacA* failed to demonstrate any signs of disease on external evaluation and disease scoring criteria (**Fig 6A, 6B**). Alterations in the corneal structure were evaluated more thoroughly with optical coherence tomography (OCT), which provides a cross-sectional image of the anterior segment in live animals for downstream and morphometric analysis. OCT revealed thickened corneal tissue and alterations in refraction for WT and complement-infected animals, indicating the presence of edema and inflammation that are characteristic of FK (**Fig 6C**). By contrast, and consistent with the external images, corneas inoculated with the Δ*hacA* mutant were indistinguishable from the sham-inoculated controls (**Fig 6D**). Tissue sections taken at day 3 p.i. were consistent with the OCT findings, demonstrating that WT and complement infected corneas were marked by ulcerated and structurally abnormal corneas, with massive immune cell infiltration in both the cornea proper and anterior chamber. Fungal hyphae were also observed through the depth of the cornea in these two groups on histology and homogenized and plated corneas from the same time point indicated similar fungal load (**Fig 6F**). Histology of the Δ*hacA* corneas, by contrast, displayed normal epithelial and stromal architecture, minimal cellular infiltration, and no visible fungal growth which was confirmed in the CFU analysis. These results, which suggest a critical role for the UPR in the establishment of corneal infection, were replicated with the full isogenic set of strains in an independent experiment using both male and female animals (**Fig S5**).

**Fig 6.**
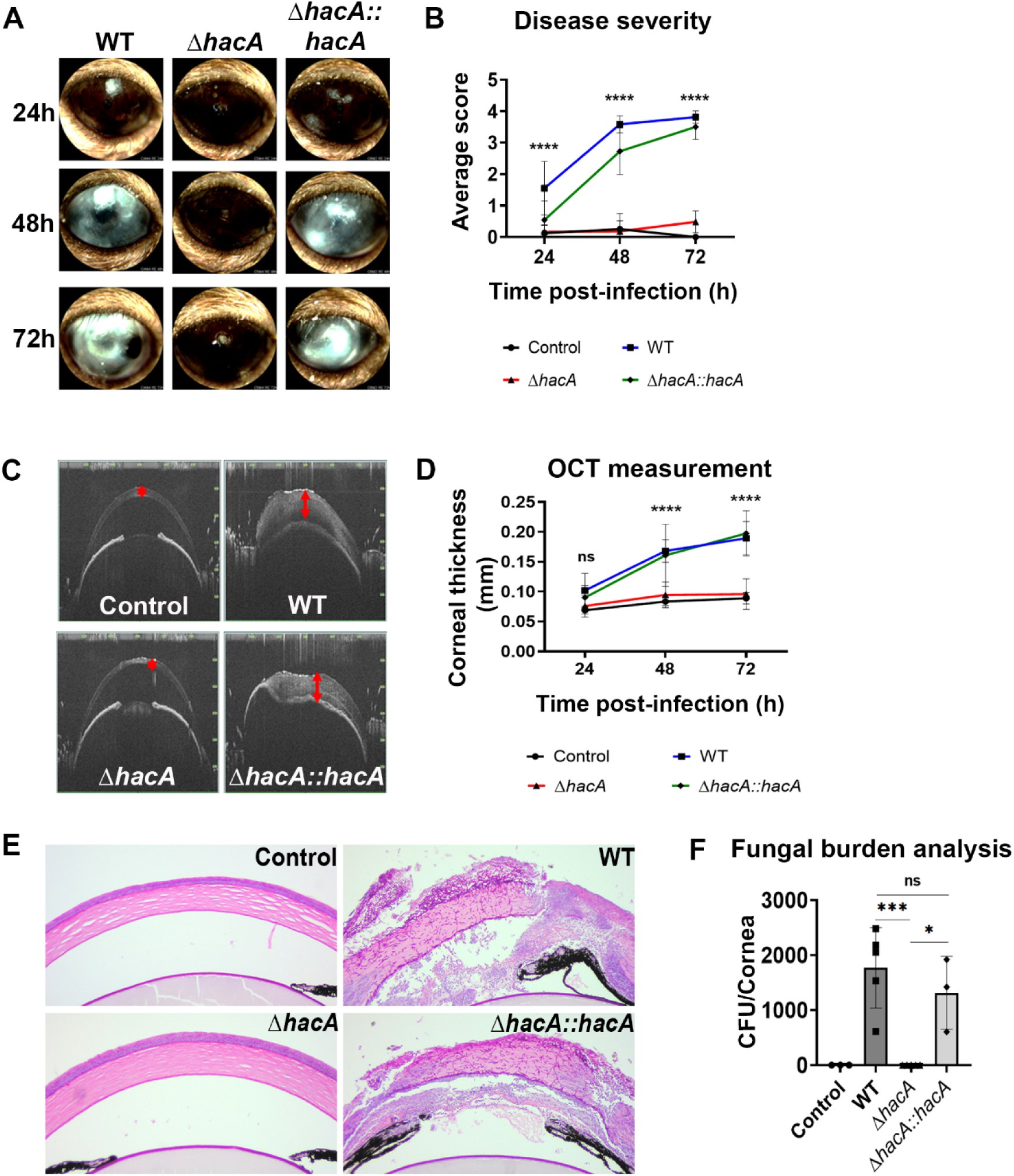
*A. fumigatus* Δ*hacA* is severely attenuated for virulence in the murine model of FK. (A) Representative external Micron IV images of infected murine corneas at 24, 48, and 72h p.i., and (B) average clinical scores (n=10/group) over the course of the infection; Data analyzed by Ordinary two-way ANOVA p-value **** <0.0001 (C) Representative cross-sectional images of the cornea from an OCT scan 72h p.i. with corneal thickness measurement (n=10/group) averaging 13 points across the cornea, (D) Corneal thickness measured across the group (n=10/group) over the course of the infection averaging 13 points across the cornea; Data analyzed by Ordinary two-way ANOVA p-value **** <0.0001 (E) Sections of cornea 72h p.i. were Periodic Acid Schiff (PASH) stained for hyphal growth and inflammatory immune cells, (F) Corneal fungal burden at 72h p.i.; Data analyzed by Ordinary one-way ANOVA p-value *** 0.0006

We next reasoned that if HacA is essential for the adaptation to the nutrient-limiting corneal environment, then the virulence of *ΔhacA* could be rescued, at least in part, through the exogenous application of glucose to the ocular surface. To test this, C57BL/6 males were infected with either the WT or *ΔhacA* strains as described above; however, a subgroup was supplemented with 50 µM glucose within the inoculum as well as through droplets applied to the corneal surface every 8 h for 3 days. PBS was applied to the corneas of the control animals in parallel. The results in **Fig 7** first show that the glucose treatments did not alter corneal clarity or structure in sham-infected animals, nor did it significantly alter the development of disease or fungal burden in the WT-infected animals. Furthermore, and contrary to our prediction, glucose did not restore any degree of disease development or fungal growth to *ΔhacA*– inoculated eyes, which at face value suggests that the primary factor driving the essentiality for HacA in establishing infection in the cornea is not a lack of a preferred carbon source. The mutant did not display hypersensitivity to other stresses likely encountered during infection, including oxidative stress or hypoxia (**Fig S4A – S4D**), altogether suggesting that the inability of *ΔhacA* to grow in the cornea is not driven by a single environmental factor, or at least one so far tested.

**Fig 7.**
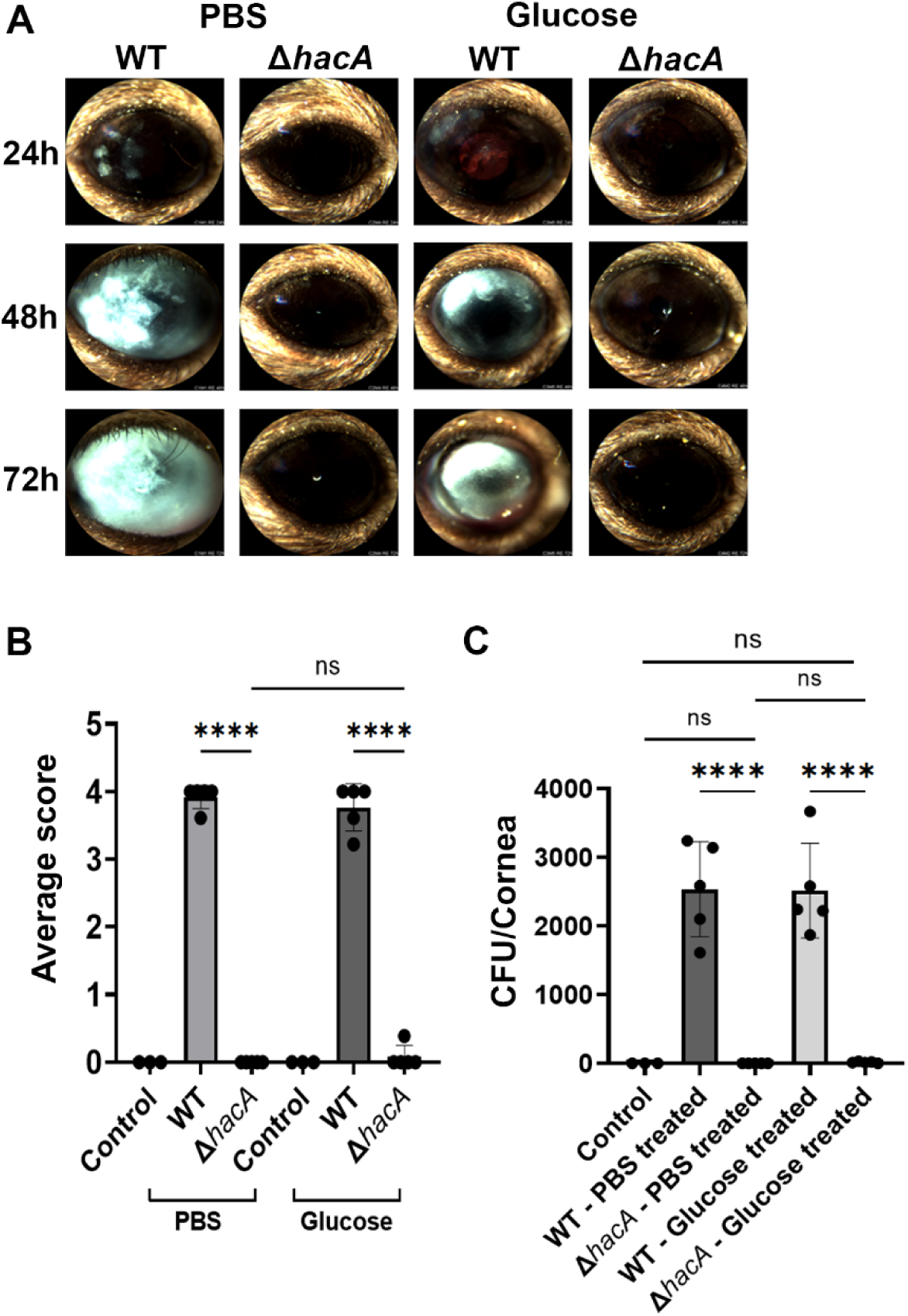
*A. fumigatus* HacA is critical for secretion. (A) Representative external Micron IV images of WT and Δ*hacA-*infected murine cornea sham (PBS) and 50 mM glucose-treated corneas every 8h over the course of the infection, (B) Average clinical scores per cornea of the sham-treated and glucose-treated WT and Δ*hacA-*infected murine (n=5/group) at 72h p.i., (C) Corneal fungal burden in the sham-treated and glucose-treated WT and Δ*hacA-*infected (n=5/group) at 72h p.i.

## DISCUSSION

Our current understanding of upstream UPR regulation in fungi, as well as its downstream influence on gene expression and biology, is largely based on studies that employ chemical stressors (e.g. DTT) that acutely misfold proteins, induce the downstream splicing of the *hacA/hac1* mRNA, and induce downstream chaperones and foldases. While these experimental parameters have indeed served to establish first principles across diverse species, it remains unclear as to whether they reflect the state of UPR functionality during infection and, by extension, if more intermediate (sub-acute) levels of UPR activation and output exist. We have used *A. fumigatus* keratitis as a platform to probe these questions, allowing us to explore a new infectious context while simultaneously leveraging prior studies that defined the *A. fumigatus* HacA-dependent transcriptomes at baseline (bUPR) and under chemically-induced ER stress (iUPR).

The UPR supports the growth of *A. fumigatus* on polymeric substrates, and while it is perhaps presumed, it has not been formally determined if nutrient stress is sufficient to promote ER stress and subsequent activation of the iUPR (14). As collagen is the primary structural component of the cornea and an ostensible source of carbon and nitrogen during infection, we utilized media containing collagen hydrolysate, gelatin, as an *in vitro* proxy for nutrient limitation. Our transcriptomic profiling of the WT revealed a broad metabolic shift towards alternative carbon utilization on gelatin, including upregulation of secreted hydrolases and membrane-bound nutrient transporters. Together with the fact that the *ΔhacA* strain displayed an enhanced growth defect on the medium, we concluded the media was indeed a source of nutrient/ER stress. Nevertheless, and in contrast to DTT-treated controls, we did not detect an increase in *hacA*^i^ abundance or chaperone induction on gelatin, indicating that a moderate shift in secretory activity is not sufficient to activate the iUPR in *A. fumigatus*. This perhaps makes sense given this saprophyte has evolved to be competitive in environments in which secretory stress is expected to be high as a rule (e.g. compost), rather than an exception. It has been postulated that this bUPR activity is important to support the endogenous levels of ER stress associated with filamentous growth, which is consistent with the finding that *ΔhacA* displays growth and developmental defects under all conditions. A steady-state activation of the UPR may also prime the fungus for rapid transitions into nutrient-poor/polymeric environments and reinforce its adaptability in the environment. It would therefore be interesting to determine if nutrient limitation has a more pronounced influence on UPR activation in non-saprophytic fungi, such as the yeast *Candida albicans*, which colonizes the gut and may experience a more stable and nutritionally replete environment at baseline.

The UPR is an important regulator of fungal pathogenicity in every organism and infectious context so far tested, including systemic candidiasis, disseminated cryptococcosis and invasive pulmonary aspergillosis (15,32–34). To our knowledge, however, there has been no direct assessment of whether the UPR of *A. fumigatus* or any other fungal pathogen is concertedly activated in host tissue. Beyond the nutritional microenvironment, it has been postulated that other stresses encountered *in vivo*, including oxidative or hypoxic stresses driven by inflammatory cells, also induce protein misfolding and cumulatively render the UPR essential for virulence (2,35,36). These stresses are known to occur during FK infection as well, as neutrophils rapidly invade the corneal tissue from the limbus and ciliary body whereupon they secrete reactive oxygen species that contribute to both fungal clearance and tissue damage (37). Indeed, deletion of the *A. fumigatus* genes encoding superoxide dismutase or the ROS-sensitive transcription factor gene, *yap1*, results in a hypersensitivity to exogenous oxidizers and reduced fungal burden in a murine model of FK (37). We have furthermore shown that murine corneas become rapidly hypoxic during FK and deletion of the hypoxia-inducible transcription factor, *srbA*, results in an avirulent phenotype (unpublished). Given the sterility of *ΔhacA*-infected corneas in our experiments, it was perhaps surprising to find a lack of iUPR induction in WT *A. fumigatus* isolated from infected and inflamed corneal tissue, both at the level of *hacA* splicing and downstream gene expression. This suggests that cumulative stresses experienced by *A. fumigatus* in the cornea do not promote the same degree of ER stress that is mediated by DTT or TM treatment *in vitro*, and that bUPR activity is sufficient to regulate growth and virulence during FK. Interesting studies would include a more thorough analysis the *A. fumigatus hacA* activation state in the lung as well as in other models of fungal infection in which the UPR is known to regulate pathogenesis.

We recognize that our assessment of UPR induction in this study is limited to and by the sensitivity of our PCR assays used to quantify *hacA* splicing and target gene expression. It, therefore, remains a possibility that more nuanced (sub-iUPR) levels of *hacA* mRNA or protein regulation exist, and these activation states would altogether fall under the current umbrella term ‘bUPR’. In support of this idea, our comparative transcriptomic data clearly indicate that HacA regulates a largely distinct subset of genes in gelatin versus GMM, despite the fact that the two conditions promote seemingly indistinguishable levels of *hacA* splicing. We, therefore, postulate that UPR output is non-binary (e.g. bUPR vs iUPR), but rather exists on a spectrum based on integrated pathway inputs, including nutrient signaling pathways, that modulate HacA transcriptional activity post-translationally. Such regulation has been observed in the mammalian system, where acetylation/de-acetylation influences the protein stability and transcriptional activity of the HacA ortholog, Xbp1 (38). Regarding *A. fumigatus*, Krishnan and colleagues revealed that ER stress leads to transcript-specific translational induction via polysome docking, which represents a form of regulation that is missed by transcriptional assays like we have employed here (19). Interestingly, *hacA* itself is not apparently regulated through this polysome mechanism, and thus it remains unclear how the proximal HacA-dependent UPR might be fine-tuned in this fungus. Ongoing studies in our laboratory include the characterization of HacA binding targets, e.g. via chromatin immunoprecipitation, under various environmental conditions. If quantitatively or qualitatively distinct binding signatures are observed, we will further interrogate HacA post-translational modifications or target-gene epigenetics as a mechanism of UPR regulation.

In order to be a successful FK pathogen, *A. fumigatus* must first establish invasive growth in the healthy (uninflamed) cornea, and then persist within the tissue in the presence of stresses imparted by the infiltrating leukocytes. Our results suggest that the *ΔhacA* mutant is unable to initiate infectious growth based on the fact that mutant-infected corneas are sterile at 72 h p.i. and do not show signs of fungal growth, altered morphology or inflammatory cells in our various clinical and pathological readouts. As we have shown that various proteolytic genes are upregulated by WT *A. fumigatus* in the cornea, it follows that the inability of *ΔhacA* to secrete such collagenases contributes, at least in part, to an inability to obtain sufficient macronutrients from the avascular and collagen-rich cornea. However, the fact that the mutant does maintain some capacity to grow on proteinaceous media *in vitro*, coupled with the inability of topical glucose drops to recover *ΔhacA* growth, together suggests that nutrient (glucose) limitation *per se* is not the primary or sole factor.

In agreement with previous reports, we have shown *ΔhacA* is hypersensitive to cell wall perturbating agents (e.g. Congo red), which likely reflects alterations in overall cell wall composition and/or architecture. Indeed, Richie et al. reported reduced glucose content in both the soluble and insoluble cell wall fractions of *ΔhacA*, suggesting a reduction in both alpha– and beta-glucan. While the generalized secretion defect of ΔhacA is thought to impact the delivery of cell wall-modulating enzymes to the hyphal apex, our transcriptomic data suggest that hacA directly or indirectly influences the expression of such genes. Among the underrepresented genes in the mutant, for example, were a myriad of endo– and exo-glucanase genes that may influence proper cell wall turnover and modification (39,40). Thus, the dysregulation of these genes may account for the generalized growth defect of the mutant in GMM, the enhanced growth defect on gelatin, and an even more marked penetration defect through the dense stromal matrix *in vivo*. Notably, one of the underrepresented genes in the mutant was *ags3*, which plays a critical role in synthesizing alpha-glucan and masking the underlying and immunogenic beta-1,3– glucan (41,42). Although we did not appreciate any signs of inflammation or disease development in *hacA*-infected corneas, it is possible that the mutant was more efficiently recognized and cleared from the cornea by resident immune cell populations or early-infiltrating neutrophils at very early time points post-inoculation.

One additional stress the fungus encounters at the ocular surface is the sequestration of iron and other essential metals by proteins in the tear film, including lactoferrin and lipocalin (37). The topical instillation of purified lactoferrin can inhibit fungal growth in the cornea, and ablation of *A. fumigatus* siderophore synthesis renders the fungus avirulent (Pearlman paper). Feng et al demonstrated that *ΔhacA* is hypersensitive to the iron chelator BPS (15), and our data indicate a down-regulation of several iron homeostasis genes in the mutant on glucose or gelatin media, including a siderophore transporter (AFUA_3G13670) and a ferric chelate reductase (AFUA_8G01310). The altered expression of metal homeostasis in the mutant is in agreement with a much larger shift in the oxidoreductase category, which supports a role for HacA in regulating mitochondrial function, redox homeostasis, and consequently optimal growth. Whether such genes are direct targets of HacA remains unclear, but it nevertheless supports an important role for the UPR in the broader metabolic adaptation to the nutrient environment.

In summary, this study is the first to evaluate the activation and influence of the *A. fumigatus* HacA transcription factor under varying degrees of ER stress. Whereas acute protein misfolding through chemical treatment can lead to detectable *hacA* mRNA splicing and chaperone induction (termed the iUPR), more nuanced levels of ER stress experienced in nutrient-limiting and corneal environments influence HacA output through another, yet-to-be-determined, mechanism. The cumulative impact of HacA on nutrient acquisition, intracellular redox balance, metal ion homeostasis, and cell wall stability, all likely contribute to the essential role of HacA for invasive growth in both the pulmonary and corneal microenvironments. This suggests that known inhibitors of the Ire1-HacA pathway in mammals may have the potential as novel antifungals in various clinical settings.

## MATERIALS AND METHODS

### Fungal strains and growth conditions

*Aspergillus fumigatus* Af293 was used as the WT organism. Fungal strains used in this study are listed in **Table S5**. The strains were maintained on a glucose minimal medium (GMM), which contains 1% glucose, Clutterbuck salts, and Hutner’s trace elements with ammonium tartrate (GMM AT) as a nitrogen source (43). To assess fungal morphology, strains were grown on yeast, peptone, and dextrose medium – YPD (2% dextrose, 2% peptone, 1%yeast extract), gelatin minimal medium (1% gelatin, Clutterbuck salts, and Hutner’s trace elements), or GMM supplemented with 1% BSA (GMM BSA) as the sole nitrogen source. For protoplasting, conidia were inoculated in GMM AT+ (GMM supplemented with 0.5% yeast extract) for 10h at 30^0^C. For protoplast recovery post-fungal transformations, GMM AT supplemented with 1.2 M sorbitol was used. The strains were grown at 35^0^C for 48h unless otherwise specified.

### Genetic manipulation

*hacA deletion*: Targeted deletion of *hacA* was performed using *in vitro*-assembled Cas9 RNPs as described earlier (30,31). Primers 685/686 were used to amplify the hygromycin phosphotransferase (*hph*) cassette from plasmid PAN 7-1 with 35 bp flanks to the *hacA* coding sequence. This repair template was introduced into Af293 protoplasts along with Cas9 RNPs designed to cut at the 5’ and 3’ ends of the *hacA* coding sequence. Transformations were performed as described previously (30,31). For selection, hygromycin B was used at a concentration of 200 µg/mL where needed. Hygromycin-resistant colonies were genotyped to confirm the loss of the *hacA* gene. All the PCRs were performed using Peltier Thermal Cycler-PTC 200 (MJ Research, Canada). The primers used in this study are listed in **Table S6**. *hacA complementation:* For complementation of the deletion mutant, a phleomycin expression cassette (*bleR* 659/660) and the *hacA* CDS were co-transformed to re-introduce *hacA* ectopically into *ΔhacA* protoplasts using the CRISPR-Cas9 technique mentioned earlier, and phleomycin at 125 µg/mL was used for selection. The presence of *hacA* was genotyped by PCR to confirm a precise integration using primers 689/690.

### Stress analyses

The indicated strains were subjected to thermal stress by spotting 2000 conidia onto the surface of a YPD plate and incubated at 35^0^ and 48^0^C for 48h. Disruption of ER homeostasis was also tested by growing the strains in liquid GMM AT in the presence of tunicamycin (TM) at 6.25 µg/mL and growth was measured at an optical density of 530nm. The mean of triplicate samples was analyzed by Brown-Forsythe and Welch’s ANOVA in GraphPad Prism 9.3.1.

For the nutritional studies, 2000 conidia were spotted onto YPD, GMM AT, GMM BSA, gelatin MM, skim milk agar and incubated at 35^0^C for 72h and colony diameter was recorded every 24h for 3 days. To test the ability of the mutant to grow on complex biological substrates, 2000 conidia were spotted on freshly isolated murine brain and lung tissue and growth was monitored over 72h. All experiments were carried out in triplicates.

For secretion stress, 2000 conidia were spotted onto GMM AT and gelatin MM supplemented with 10 µg/mL of brefeldin A (BFA). All experiments were carried out in triplicates and growth was recorded after incubation for 72 h at 35 °C.

For oxidative stress analysis, WT and Δ*hacA* were inoculated in GMM AT at 10^6^ conidia/mL concentration in 96-well plates with and without agents for inducing oxidative stress (0.5–8 µM H_2_O_2_ or 5-60 µM menadione). Oxidative stress by H_2_O_2_ was also tested on GMM AT agar plates by spotting 2000 conidia using similar conditions as mentioned above. The mean of triplicate samples was analyzed Unpaired T-test in GraphPad Prism 9.3.1.

For hypoxic stress analysis, 2000 conidia were spotted on GMM plates in normoxia (∼21% O_2_) or hypoxia (1% O_2_, 5% CO_2_). The experiment was carried out in triplicates and growth was recorded after incubation for 72 h at 35 °C.

To test susceptibility to Congo red, 2 µl of a 1.0 × 10^6^ conidial suspension was spotted onto GMM medium containing the indicated concentrations of Congo red. All pictures shown were taken after 72h incubation at 35°C.

### Azocollagen assay

Secretion of collagenase in *A. fumigatus* was quantified using Azocoll (Millipore Sigma) hydrolysis as previously described (44). Conidia were inoculated at a concentration of 10^5^/mL in 25ml GMM supplemented with 10% heat-inactivated fetal bovine serum (ATCC, USA) as shown previously (14). The cultures were grown at 200 rpm for 72 h at 35^0^C. The supernatant was removed from the cultures and centrifuged at 12,000rpm for 5 mins and a 100uL aliquot was added to 400uL of pre-washed Azocoll (5mg/mL) prepared using a buffer containing 50 mM Tris-HCl (pH 7.5), 0.01% (wt/vol) sodium azide and 1mM CaCl_2_ (44). The samples were then incubated at 37^0^C for 3h with gentle shaking. The tubes were centrifuged at 10,000 rpm for 3min, and the absorbance was measured at 520 nm to measure the release of azo dye from the supernatant. These values were normalized to the dry weight (g) of the different strains using the 72h biomass. A mean of four replicate samples was analyzed by Brown-Forsythe and Welch’s ANOVA in GraphPad Prism 9.3.1.

### Murine model of keratitis

*Ethical Statement*: All animal studies were performed in accordance with the Association for Research in Vision and Ophthalmology (ARVO) guidelines for the use of animals in vision research and were approved by the University of Oklahoma Health Sciences Center Institutional Animal Care and Use Committee (Protocol: 20-060-CI).

#### Inoculum

The indicated strains were grown on solid GMM AT, under selection where relevant, for 48h at 35^0^C. Conidia were collected by washing the surface with phosphate-buffered saline (PBS) and filtering it through miracloth before washing twice with PBS. 5×10^6^ conidia were grown in YPD for 4h at 35^0^C till they reached the stage where most of the conidia are swollen. At this stage, the culture was washed and resuspended in 500uL of PBS. The strains were normalized at OD 360nm and 5uL of this suspension was then used to infect the eyes.

#### Mice

6-8-week-old C57B6/6J mice (Jackson Laboratory) were used for the animal experiments. A corticosteroid-epithelial abrasion model was used to carry out the infection studies. Animals were immunosuppressed with an intraperitoneal (IP) injection of 100 mg/kg Depo-Medrol (Zoetis, USA) on the day preceding inoculation (day –1). On day 0, mice were anesthetized with 100 mg/kg ketamine and 6.6 mg/kg xylazine IP and an Algerbrush II was used to abrade the central epithelium of the right eye to a diameter of 1 mm. 5uL of WT, Δ*hacA* and Δ*hacA*::*hacA* inocula (described above) was applied to ulcerated eyes and removed with a Kim wipe after 20 minutes. A single dose of Buprenorphine SR (1 mg/kg) was administered subcutaneously for analgesia. The contralateral eye of each animal remained uninfected in accordance with the ARVO ethical guidelines.

#### Micron IV slit-lamp imaging

The animals were monitored every day p.i. by capturing live corneal photographs obtained from mice anesthetized with isoflurane with a Micron IV slit-lamp imaging system (Phoenix Research Labs Inc., CA, USA) for up to 72h p.i.

#### Anterior segment spectral-domain optical coherent tomography (OCT)

The corneal thickness was measured for all the mice at 48 and 72h p.i. using an OCT equipment (Leica Microsystems, IL, USA). Before imaging, the mice were anesthetized using isoflurane and a 12 mm telecentric lens was used to generate a 4×4 mm image. The reference arm was calibrated and set to 885 by the manufacturer. The InVivoVue driver software was used to further analyze the images. To quantify the corneal thickness, measurement was performed using an 11×11 spider plot to cover the entire eye, and an average of 13 readings were taken, and analyzed by Ordinary one-way ANOVA in GraphPad Prism 9.3.1. Algerbrushed sham-infected eyes were used as a control.

#### Histopathological examination

Control and infected eyes were harvested and fixed with 10% neutral buffered formalin for 4 h followed by 70% ethanol until further processing. The eyes were sectioned (5 μm thick) and stained with Periodic acid Schiff-hematoxylin (PASH) to evaluate the fungal cell wall and the host inflammatory response in the murine model of FK.

#### Fungal burden determination

Corneas were dissected aseptically and homogenized in 1 ml of 2mg/ml collagenase buffer, and 100 μl aliquot was plated in triplicate onto inhibitory mold agar (IMA) plates and incubated at 35°C for 24 h to determine the number of colony-forming units (CFU) per cornea. The statistical analysis on the colony counts was performed across the group using Ordinary one-way ANOVA in GraphPad Prism 9.3.1. The sham-infected eyes were used as controls.

#### Clinical scoring

The micron images were visually clinically scored using the criteria previously established for keratitis by two blinded reviewers (45). The eyes were graded on a scale of 0 to 4 and averaged based on surface regularity, area and density of opacity. The statistical analysis on the average score per cornea was measured across the group using Ordinary one-way ANOVA in GraphPad Prism 9.3.1.

### Glucose supplementation *in vivo*

The murine model described earlier was used to test the rescue of fungal growth *in vivo* using glucose. Mice were infected with WT and Δ*hacA.* The inoculum was prepared as described earlier and glucose was added to it at a final concentration of 50mM. The treatment regimen included the topical application of 50 mM glucose to the eyes every 8 hours i.e. 3 times/day from the time of the infection. The endpoint assessment was carried out as described earlier for blinded clinical scoring and eyes were harvested at 72 h p.i. for fungal burden analysis.

### qRT-PCR and RNA-sequencing

RNA was extracted using Qiagen RNeasy Mini Kit (Maryland, USA) according to the manufacturer’s protocol and DNase-treated using the DNase I kit (Millipore Sigma, Massachusetts, USA). The quantity and quality of RNA were determined using the Nanodrop 2000 (Thermo Fisher Scientific, Massachusetts, USA). The RNA was normalized for cDNA conversion using ProtoScript® II First Strand cDNA Synthesis Kit (New England Biolabs, Massachusetts, USA) following the manufacturer’s protocol. qRT-PCR was performed using Luna Universal qPCR master mix (SYBR green; NEB, USA) on the QuantStudio™ 3 Real-Time PCR System (Thermo Fisher Scientific, Massachusetts, USA) and analysis was performed using QuantStudio™ Design and Analysis Software v1.5.2. The fold-expression changes were calculated using the 2^-ΔΔCt^ method and followed by analysis using MS-Excel and Graph Pad Prism 9.3.1.

#### Protease expression

Murine cornea were infected using the corticosteroid-epithelial abrasion model with WT *A. fumigatus* and the expression of proteases with collagenase activity – *alp1* (761/762), *mep* (763/764), *dppIV* (765/766) and *dppV* (767/768) was studied at 48 h p.i. The infected eyes were harvested following euthanasia under an IACUC-approved procedure followed by corneal dissection. Green bead lysis kits (Next Advance, New York, USA) that contain stainless steel beads in combination with TissueLyser LT (Qiagen, Maryland, USA) at 50 oscillations/s for two 30 s cycles were used to homogenize the corneal tissue followed by RNA extraction. The expression was normalized to β-tubulin which was used as the housekeeping gene. All qRT-PCR experiments were performed at least in duplicates.

#### RNA sequencing

8ml of GMM AT and gelatin MM were inoculated with WT or Δ*hacA* with a density of 1.0 x 10^5^/mL in 6-well plates and incubated in 35^0^ C static culture for 48 h. As a positive ER control, GMM cultures were treated with 10 mM DTT 2 h prior to harvest. After 48 h, the fungal biomass was harvested and placed in a 1.5 mL microcentrifuge tube with 1.0 mm diameter glass beads (BioSpec, Bartlesville, OK, USA) and tissue was homogenized using TissueLyser LT (Qiagen, Maryland, USA) same as with the corneal tissue mentioned previously. RNA extraction was perfomed as described earlier. Briefly, the quality of the RNA was checked using Bioanalyzer 2100 (Agilent, CA, USA), and concentrations from NanoDrop (Thermo Fisher Scientific, MA, USA) readings were used. Stranded RNA-seq libraries were constructed using NEBNext poly(A) mRNA isolation kit along with the SWIFT RNA Library Kit (NEB, MA, USA) according to the established protocols at Laboratory for Molecular Biology and Cytometry Research (LMBCR) at The University of Oklahoma Health Sciences Center (OK, USA). The library construction was done using 100ng of RNA. Each of the 18 libraries was indexed during library construction in order to multiplex for sequencing. Libraries were quantified using Qubit 4 fluorometer (Invitrogen, MA, USA) and checked for size and quality on Bioanalyzer 2100 (Agilent, CA, USA). Samples were normalized and pooled onto a 150 paired end run on NovaSeq (Illumina, CA, USA) to obtain 50M reads per sample. Differentially expressed genes (DEGs) between groups were considered to be significant if a log2 fold change (FC) was at least ≥4-fold. Gene ontology (GO) and Functional Catalogue (FunCat) were used to systemically categorize the DEGs. A few select genes were validated using qRT-PCR which was performed at least in duplicates.

### *hacA* splicing

To qualitatively detect the presence of the two forms (uninduced form *hacA^u^* – 665 bp; induced form *hacA^i^* – 645 bp) in the WT-infected corneas at 24 and 48h p.i., RT-PCR was performed using 689/690 and run on a 3% agarose gel. Cultures of *A. fumigatus* grown for 48h in liquid GMM AT, gelatin MM, and GMM AT treated with 10 mM DTT for 2 h before extraction of total RNA were used as controls. For expression analysis, qRT-PCR with primer pairs that yielded PCR products around the predicted *hacA* unconventional intron were used to amplify the two forms (uninduced form *hacA^u^* – 749/750; induced form *hacA^i^* – 749/751) of the gene coding for HacA in *A. fumigatus* (Afu3g04070). Primer pairs 752/753 and 754/755 were used to quantify the expression of the UPR target genes – *bipA* and *pdiA* respectively.

## Acknowledgements

This work was supported by the National Institutes of Health (P20GM134973, R01EY021725 and P30EY021725) and Research to Prevent Blindness (Career Development Award to KKF and an unrestricted grant to Dean McGee Eye Institute). We thank Mark Dittmar and staff (DMEI Animal Research Facility), Linda Boone (DMEI Imaging Core), and the Institutional Research Core Facility at OUHSC for the use of the Core Facility which provided total RNA library construction and sequencing.

